# Long-term iron deficiency and iron supplementation exacerbate acute DSS-induced colitis and are associated with significant dysbiosis

**DOI:** 10.1101/585174

**Authors:** Awad Mahalhal, Michael D. Burkitt, Carrie A. Duckworth, Georgina L. Hold, Barry J. Campbell, D. Mark Pritchard, Chris S. Probert

## Abstract

Patients taking oral iron supplementation often suffer from gastrointestinal side effects. We have previously shown that acute alterations in oral iron exacerbate dextran sodium sulphate (DSS) induced colitis and are associated with dysbiosis. As patients take iron supplementation for long periods, we asked whether this too would influence colitis and the microbiome. We assessed the impact of long-term changes in dietary iron, by feeding chow containing 100ppm, 200ppm and 400ppm (reflecting a deficient, normal or supplemented diet, respectively) for up to 9 weeks to female wild-type C57BL/6 (WT) mice in presence or absence of chronic colitis, or acute colitis induced after 8 weeks, induced by DSS. Assessment was made based on (i) clinical and histological severity of colitis, and (ii) faecal microbial diversity, as assessed by sequencing the V4 region of *16S* rRNA. In mice with long term changes to their dietary iron, reduced iron intake (100ppm iron diet) was associated with increased weight loss and histology scoring in the acute colitis model. Chronic colitis was not influenced by altering dietary iron however there was a clear change in the faecal microbiome in the 100 and 400ppm iron DSS-treated groups and in controls consuming the 400ppm iron diet. Proteobacteria levels increased significantly at day-63 compared to baseline and Bacteroidetes levels decreased in the 400ppm iron DSS group at day-63 compared to baseline; mirroring our previously published work in acute colitis. Long term dietary iron alterations clearly affects gut microbiota signatures but do not appear to exacerbate chronic colitis. However, acute colitis is exacerbated by changes in dietary iron. More work is needed to understand the impact of iron supplementation of the pathologenesis of IBD and rise that possiblity that the change in the microbiome, in patients with colitis, is a consequence of the increase in luminal iron and not simply the presence of colitis.

## Introduction

Inflammatory bowel disease (IBD) is a debilitating, relapsing-remitting long-term condition of the gastrointestinal tract that affects around 240,000 people in the UK (1) (2, 3). Approximately one-third of patients develop iron deficiency anaemia because of intestinal bleeding and/or malabsorption (4-7). Iron deficiency (ID) may be treated effectively by intravenous or oral iron replacement (8). These therapeutic options have different side effect profiles (9), and may have other off target effects e.g. iron is a growth factor for some bacteria (10). Unabsorbed oral iron supplements and gastrointestinal bleeding result in an increase in luminal iron which may exacerbate IBD and lead to increased proliferation and virulence of some bacteria (11-13). Intestinal bacterial dysbioses have been associated with relapse of IBD (14, 15). It is not clear whether relapsing inflammation leads to dysbiosis by modulating luminal iron (16).

Chronic inflammation of the intestinal tract is the main feature of IBD. Intestinal epithelial cells (IECs) provide a single superficial layer on the intestinal mucosa and act as the first defensive barrier against the luminal content of the gut and protector of the underlying tissues. IECs have important roles, secreting antimicrobial substances [defensins] and communicating with intestinal immune cells using soluble mediators, chemokines and cytokines (17, 18). There is mounting evidence that alterations in immune regulatory pathways, including inflammasome activation pathways drive changes in gut microbiotal diversity (19). The mucosal barrier not only defends against luminal pathogens, but also actively shapes the peri-mucosal niche, thereby regulating the composition of the mucosa-associated microbiota (20).

Based on this evidence, we hypothesised that iron supplementation (and or bleeding) in IBD patients could change the composition of the gut microbiota and potentially influence the natural history of IBD. To investigate this, we assessed the long-term effects of altering dietary iron consumption on intestinal microbiota in murine models of colitis to eliminate any confounding factors based on background genetics that would be inevitable in a human population.

## Materials and Methods

### Animals

Wild type C57BL/6 female mice, aged 8-9 weeks old, were purchased from Charles River Laboratories (Margate, UK). Six groups of 8 mice were studied: three control groups and three DSS-treated groups, all of which were maintained for 63-days. Mice received standard chow and water *ad libitum*, during an acclimatisation period of at least one week. Animals were then individually caged in a room with controlled temperature, humidity and a pre-set dark-light cycle (12 h: 12 h) in a specific pathogen-free animal facility. For each group of experiments, mice were matched for age and body weight. The care of, and experimentation on, mice was carried out in accordance with UK Home Office regulations (project licence no: 70/8457) and the project was reviewed by the University of Liverpool Animal Welfare and Ethical Review Body (AWERB).

### Diets

When eating a normal (standard) iron diet, mice were fed Rat and Mouse Breeder and Grower Pelleted CRM (P) chow (Special Diets Services (SDS), Witham, Essex, UK) which contained 200 part per million (ppm) iron in 10mm compression pellets. Two modifications of this standard iron diet were also used: the first was CRM (P) 100ppm iron (Fe) diet where the CRM (P) formulation was used with reduced iron content (0.01% Fe), this was called the half standard iron diet (100ppm iron). The second modification was the CRM (P) 400ppm iron diet: again the CRM (P) formulation was used, but the iron content was increased (0.04% Fe), this was called the double standard iron diet (400ppm iron).

### Controls

Mice in the three control groups received the standard, half standard or double standard iron diets respectively for 63 days. After 53-days, each group was divided into two, half carried on as controls and half were treated with 2% DSS as described below (Supplementary Figure 1).

### Induction of chronic colitis using dextran sodium sulphate (DSS)

Three groups of 8 mice (taking standard, half standard and double standard iron diets respectively) were given a 1.25% solution of dextran sulfate sodium (M.W. 36,000 – 50,000Da; Catalogue number: 160110; Lot number: 6683K; MP Biomedicals, LLC, UK) in their drinking water for 5-days to induce colitis (Supplementary Figure 1). Mice were allowed to recover for 16 days and then the DSS-treatment was repeated for a total of three cycles (21).

### Induction of acute colitis using DSS

Three groups of 4 mice which had been on the diets for 53 days (control groups) received 2% DSS for 5-days in drinking water, followed by 5-days of plain drinking water, to induce acute colitis. All mice were euthanised on day-63.

### Histopathological scoring of colonic inflammation

The distal colon was removed, fixed in 4% neutral buffered formalin, dehydrated, wax-embedded and then cut into 4μm sections. The sections were stained with haematoxylin and eosin (H&E). Inflammation was reported using the inflammatory scoring system described by Bauer *et al.* (22). Fibrosis was assessed using Masson’s trichrome staining (NovaUltra™ Masson’s Trichrome Stain Kit (Fisher Scientific UK Ltd)) (23). A researcher blinded to the treatment group assessed all slides.

### Assessing the degree of gut inflammation by measuring faecal calprotectin concentrations

Faecal calprotectin concentration was measured using the S100A8/S100A9 ELISA kit (Immundiagnostik AG, Stubenwald-Allee 8a, Bensheim, Germany) from faecal samples collected from each mouse, on day-1, 21, 42 and 63 in the chronic colitis study, and on day-1 and day-10 of the acute colitis study in control mice.

### Faecal iron concentration

The faecal iron (Fe^2+^ and Fe^3+^) concentration was measured using an iron immunoassay kit (MAK025, Sigma-Aldrich) from the same faecal pellets that were collected for calprotectin assessment.

### Faecal bacterial DNA extraction and sequencing

2g of faeces was used for bacterial DNA extraction using Stratec Kit (PSP® Spin Stool DNA Plus Kit, STRATEC Molecular GmbH, D-13125 Berlin) following the supplier’s protocol. The extracted DNA was sent to the Centre for Genomic Research at the University of Liverpool to undertake the rest of amplicon library protocol 16S [Metagenomic Sequencing Library]. Primers described by Caporaso *et al. (24)* were used to amplify the V4 region of 16S rDNA F: 5’ACACTCTTTCCCTACACGACGCTCTTCCGATCTNNNNNGTGCCAGCMGCCGCGGTAA 3’ and R: 5’GTGACTGGAGTTCAGACGTGTGCTCTTCCGATCTGG ACTACHVGGGTWTCTAAT3’.

Approximately 5 µg of extracted DNA was used for first round PCR with conditions of 20 sec at 95°C, 15 secs at 65°C, 30 sec at 70°C for 10 cycles then a 5 min extension at 72°C. Samples were purified using Axygen SPRI Beads. The second-round PCR was performed to incorporate Illumina sequencing adapter sequences: 15 cycles of PCR were performed using the same conditions. Samples were re-purified then quantified using Qubit and assessed using the Fragment Analyser. Successfully-generated amplicon libraries were sequenced (25).

The final libraries were pooled in equimolar amounts using the Qubit and Fragment Analyser data and 350-550 bp size-selected on the Pippin Prep. The quantity and quality of each pool were assessed by Bioanalyzer and subsequently by qPCR using the Illumina Library Quantification Kit from Kapa on a Roche Light Cycler LC480II according to manufacturer’s instructions. The pool of libraries was sequenced on one lane of the MiSeq at 2×250 bp paired-end sequencing. To help balance the complexity of the amplicon library 15%, PhiX was spiked in (25).

### Bioinformatics analysis

Initial processing and quality assessment of the sequence data was performed using an in-house pipeline. Base-calling and de-multiplexing of indexed reads were conducted by CASAVA version 1.8.2 (Illumina). The raw fastq files were trimmed to remove Illumina adapter sequences where any reads that matched the adapter sequence over at least three bp was trimmed off. The reads were further trimmed to remove low-quality bases (reads <10 bp were removed). Read pairs were aligned to produce a single sequence for each read pair that would entirely span the amplicon. Sequences with lengths outside the expected range were excluded (25). The sequences passing the above filters for each sample were pooled into a single file. A metadata file was created to describe each sample. These two files were analysed using Qiime, version 1.8.0 (Caporaso *et al*., 2010) (26). Similar sequences were clustered into groups, to define OTUs of 97% similarity. OTU-picking was performed using USEARCH7 (Edgar *et al.*, 2010) (27). The Greengenes database version 12.8 (McDonald *et al.*, 2012) (28), was used for reference-based chimaera detection (25). OTU tables were repeatedly sub-sampled (rarefied). For each rarefied OTU table, three measures of alpha diversity were estimated: chao1, the observed number of species, and the phylogenetic distance. For inter-sample comparisons (beta-diversity), all datasets were rarefied, and tables were used to calculate weighted and unweighted pair-wise UniFrac matrices using Qiime. UniFrac matrices were then used to generate UPGMA (Unweighted Pair-Group Method with Arithmetic mean) trees and 2D principal coordinates plots (PCoA).(25)

### Statistics

Normally distributed physiological and biochemical data were assessed by analysis of variance followed by multiple comparisons Dunn’s test and non-normally distributed data have been evaluated by Kruskal-Wallis test followed by multiple comparisons Dunn’s test (Stats Direct version 3.0.171). For the bioinformatic analysis of microbiota data, Kruskal-Wallis H-test was used with the false discovery rate (FDR) Storey’s (multiple correction tests). The q-value is the adjusted p-value based on FDR calculation, where statistical significance was declared at p<0.05.

## Results

### Chronic DSS-induced colitis induces C57BL/6 weight loss

Colitis was reproducibly induced by 1.25% DSS. All mice lost body weight from day-6 and maximal weight loss occurred at day-8 of each cycle. Mice receiving the 100ppm iron diet appeared to lose more weight than other groups, but this difference was not significant (Fig. 1-a). All control mice, irrespective of the iron dosing, showed a steady increase in body weight. However, mice fed 400ppm iron diet showed a significant weight gain during the whole 63 days this reflect the nutritional factor effect (Supplementary Figure 2).

**Figure 1-a:**
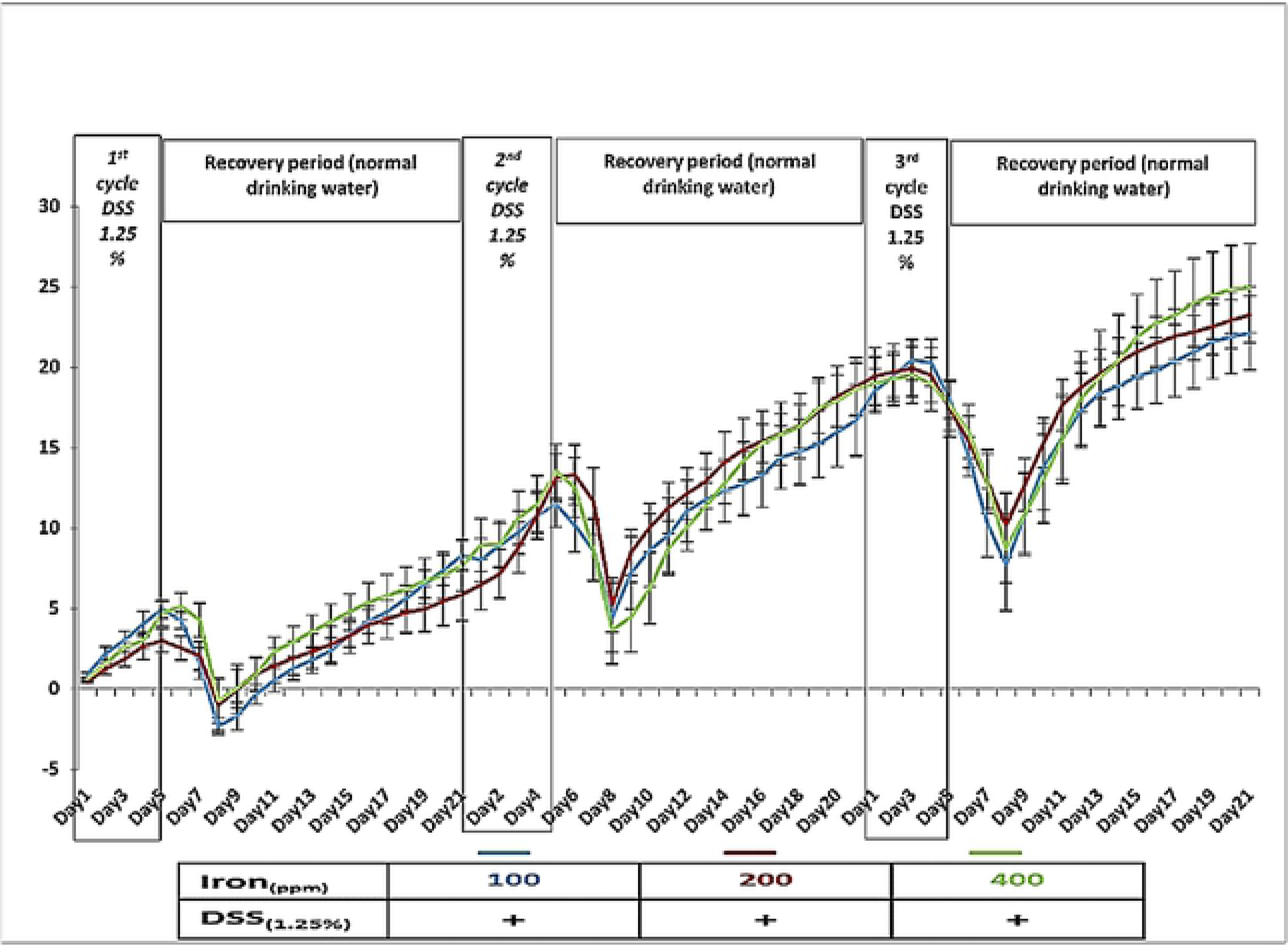
Percentage of weight change in mice (100ppm iron (blue), 200ppm iron (red) and 400ppm iron (green)) during three cycles of 1.25% dextran sulphate sodium-induced colitis during the 63-day period. Data are presented as a mean ± standard error of the mean. Statistical differences were assessed by the Kruskal–Wallis test followed by Dunn’s multiple comparison tests. (n=8 female mice per group).

### Acute DSS-colitis induced weight loss is more severe in mice fed 100ppm iron diets

Acute DSS colitis was induced after 53 days of dietary manipulation in a subset of mice that had consumed different amounts of dietary iron during this time: all developed colitis. Weight loss began earlier (day-3) in the 100ppm iron group than in the 200 and 400ppm iron DSS-treated groups (Fig. 1-b). During this acute DSS cycle, mice fed 100ppm iron lost significantly (P<0.001) more weight than the other treated groups.

**Figure 1-b:**
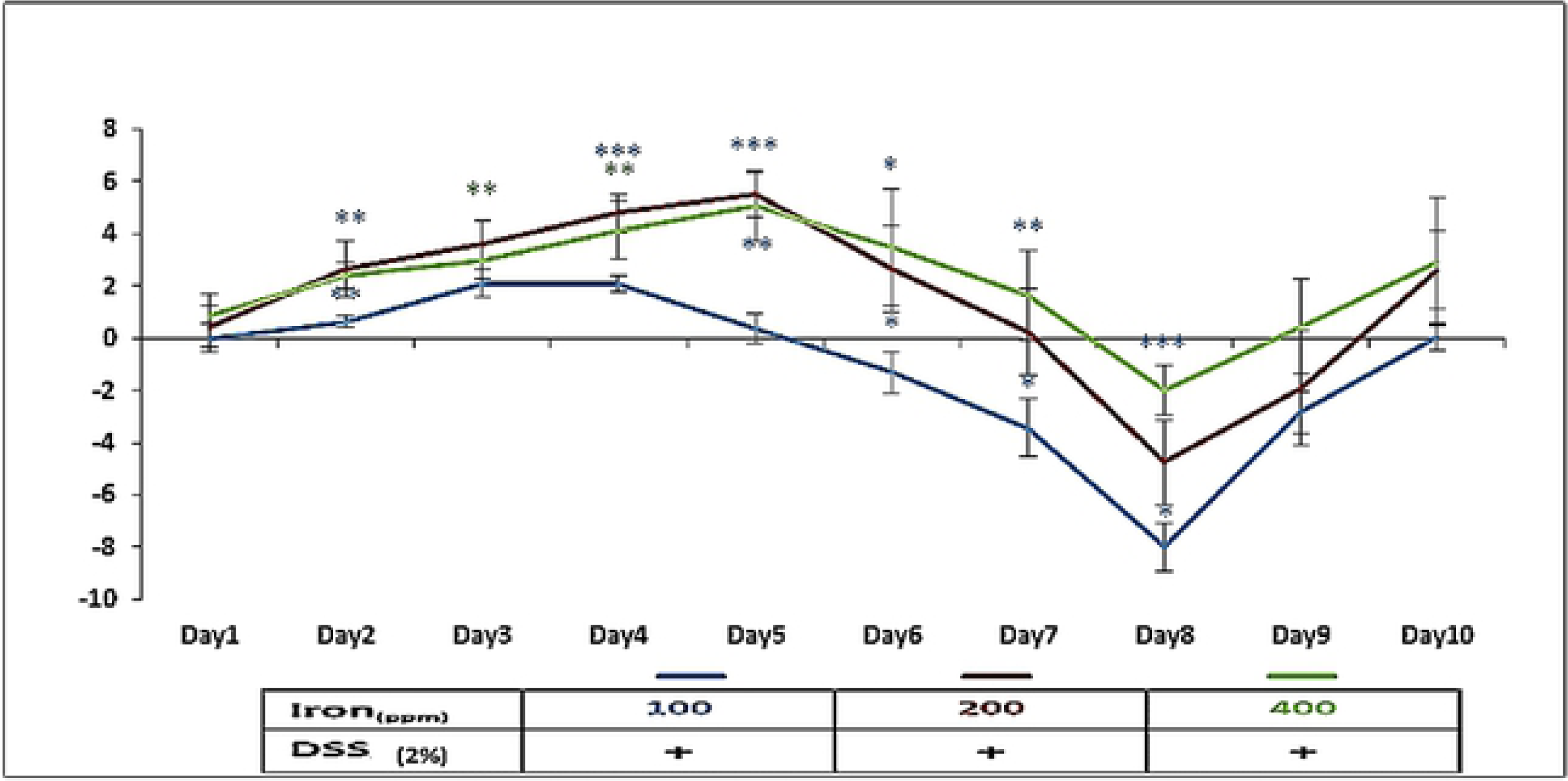
Percentage of weight change in mice (100ppm iron (blue), 200ppm iron (red) and 400ppm iron (green)) during 2% dextran sulphate sodium-induced colitis. Data are presented as a mean ± standard error of the mean. Statistical differences were assessed by the Kruskal–Wallis test followed by Dunn’s multiple comparison tests, compared with standard chow group. (n=4 female mice per group). * P<0.05, ** P<0.01, *** P<0.001.

### Histopathological changes caused by acute and chronic DSS treatment

At autopsy all mice that had been treated with repeated cycles of 1.25% DSS showed histological evidence of mild chronic colitis (Fig. 2-a; I, II and III), and those receiving acute DSS treatment had moderately severe acute colitis (Fig. 2-a; VII, VIII and IX). By contrast the colons of control untreated mice appeared histologically normal (Fig. 2-a; IV, V and VI). The colitis scores were significantly greater (P<0.01) in the mice that had been treated with 2% DSS after consuming either 100 or 400ppm iron, compared with the mice that had received 200ppm iron and all the mice that received cycles of 1.25% DSS (Fig. 2-b).

**Figure 2-a:**
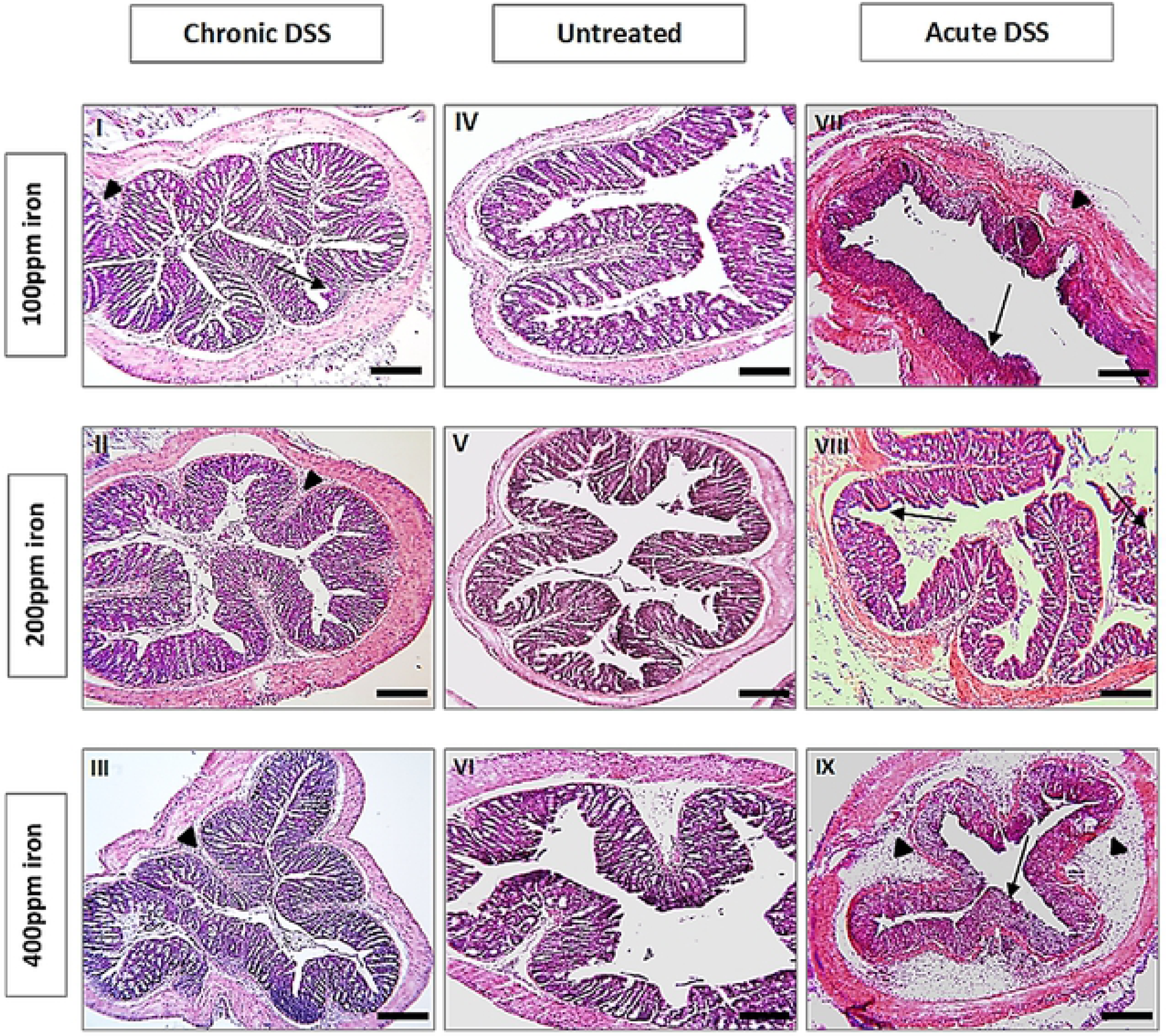
Illustrative H&E-stained segments of distal colon from untreated (n=4), 1.25% (n=8) and 2% DSS-treated mice (n=4). Mice received either water (control) (IV, V, VI), 1.25% DSS for 5 days and full recovery period 16 days on normal water (I, II, III) or 2% DSS for 5 days and followed by another 5 days on plain drinking water before they were euthanised (VII, VIII, IX). Arrowheads highlight submucosal oedema; arrows highlight almost complete loss of colonic epithelium. Scale Bar: 100 µm.

**Figure 2-b:**
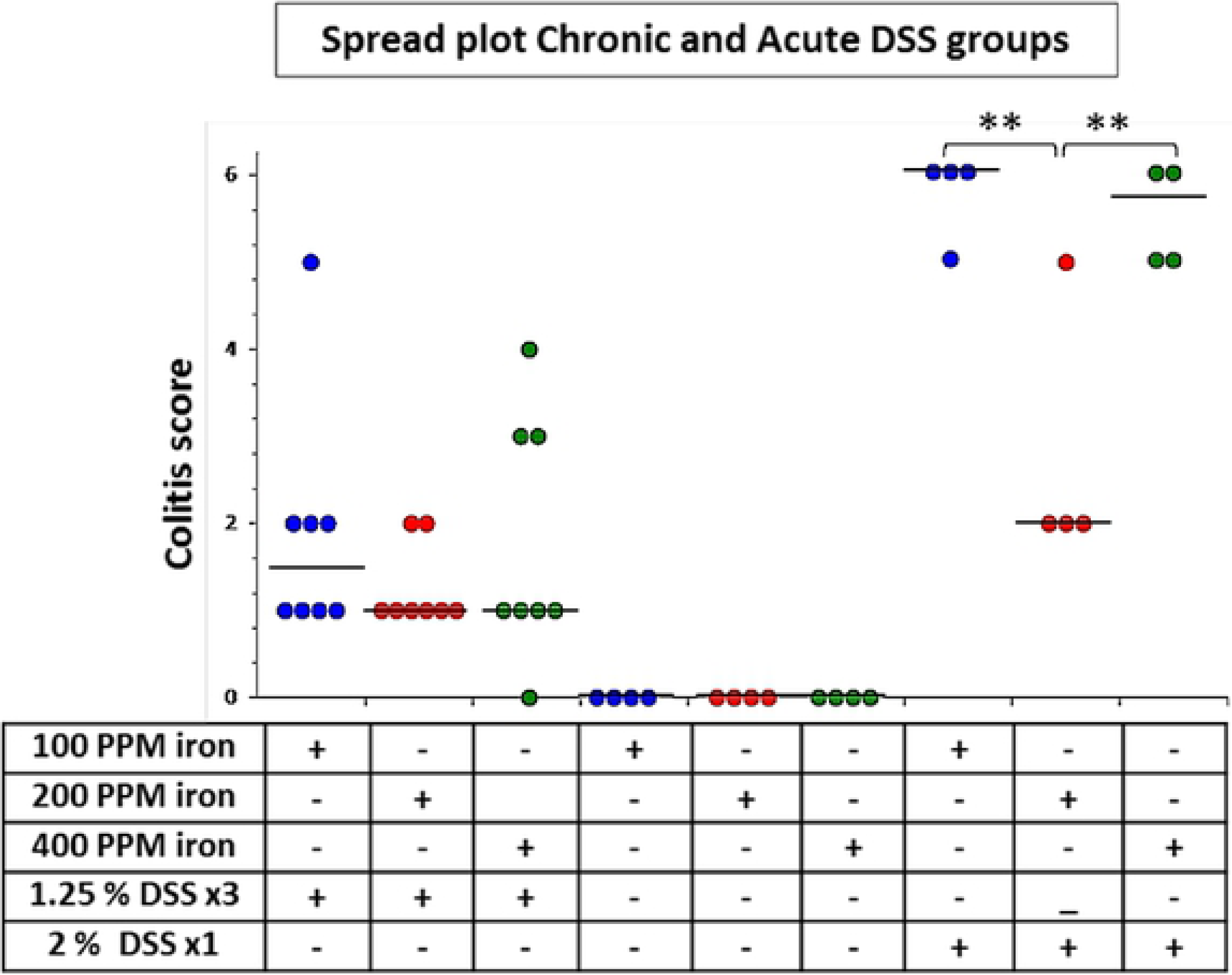
Inflammation (colitis) scores for all groups’ DSS-treated (n=8 (63-days) and n=4 (10-days) mice per group) and untreated (controls) mice on different iron diets n=4 per group (63-days). Horizontal lines at the median. Differences tested by One-way ANOVA followed by multiple comparisons Dunn’s test. **P<0.01.

### Analysis of intestinal fibrosis in chronic colitis in mice treated with repeated cycles of dextran sulphate sodium

Masson’s trichrome staining was used to assess the degree of fibrosis following chronic DSS treatment (Supplementary Figure 3). Mice in the 100ppm iron DSS-treated group had significantly more fibrosis (P<0.05) than the DSS-treated mice receiving 200ppm iron and 400ppm iron diets (Supplementary Figure 4).

### Faecal calprotectin concentration in chronic and acute DSS-treated mice

Faecal calprotectin concentrations appeared to increase after each cycle of 1.25% DSS treatment in mice consuming the 400ppm iron diet; the differences were statistically significant (P<0.01) between day-21 and day-63 (Supplementary Figure 5): this was not seen in other mice. Thus, mice with double standard iron diet appeared to develop more inflammation at molecular level by assessment of faecal calprotectin concentration.

For the acute DSS experiment, faecal calprotectin concentration increased significantly in each DSS-treated group. The change in faecal calprotectin was greater in the 100ppm iron diet DSS-treated mice than in the other groups (Supplementary Figure 6). Thus, mice consuming half-standard iron diets also appeared to develop more molecular inflammation after acute colitis induced.

### Faecal iron concentrations

In the chronic colitis experiment, DSS-treated mice consuming 400ppm iron showed a difference in faecal iron concentration between day-1 and day-63 only. Mice in the 100 and 200ppm treated groups that received DSS both showed significant differences at day-1 vs day-21, 42 and 63 (Fig. 3-a) consistent with the presence of luminal iron from bleeding resulting from colitis. Faecal iron concentration increased significantly in control mice (63 days on diet alone) taking 200 and 400ppm diets, but did not change with time in those mice consuming 100ppm iron (Fig. 3-a).

**Figure 3-a:**
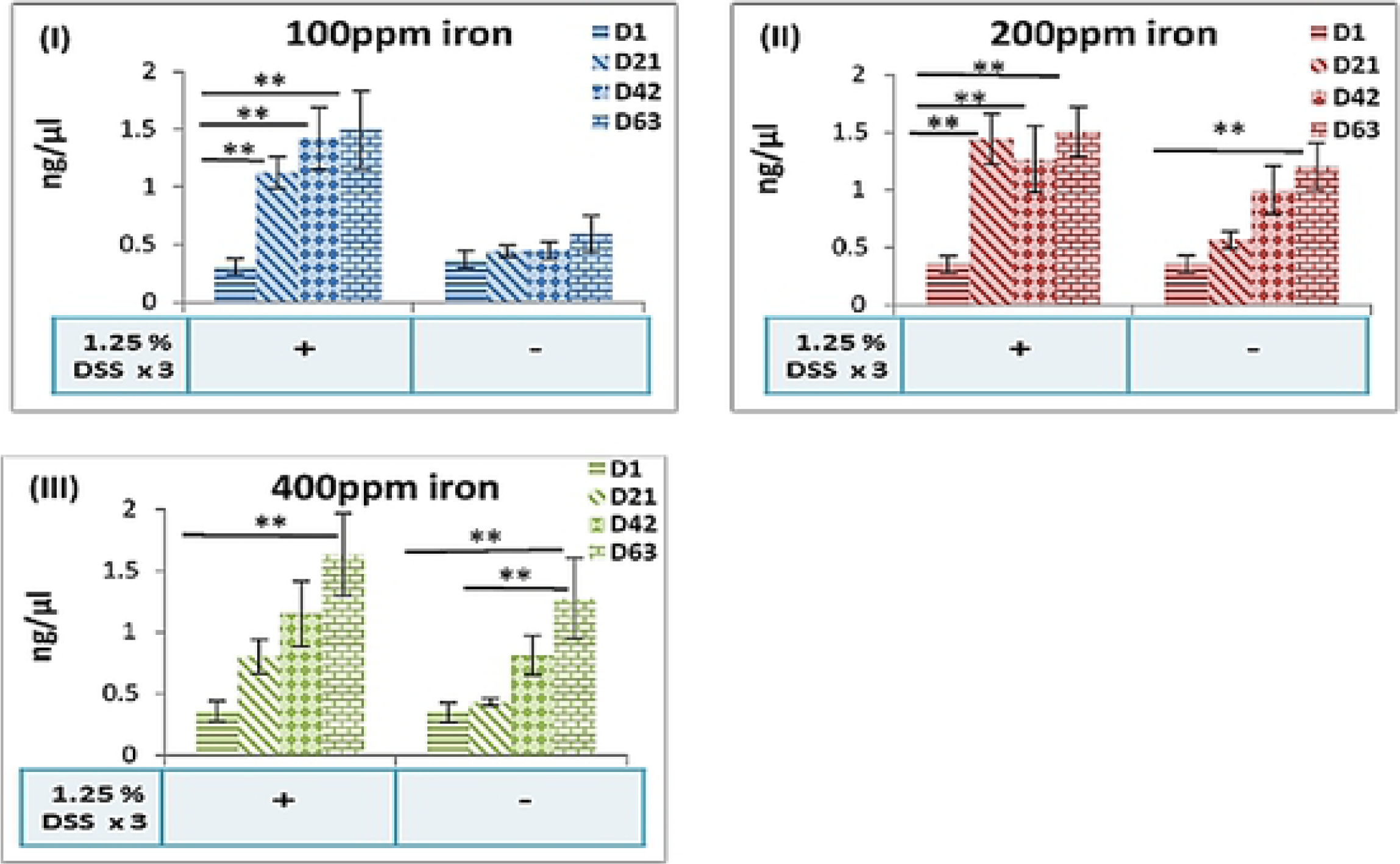
Faecal iron concentration at four different time points day-1, 21, 42 and 63 separately. (I) Faecal iron in 100ppm iron DSS-treated and untreated groups (II) faecal iron in 200ppm iron DSS-treated and untreated groups (III) faecal iron in 400ppm iron DSS-treated and untreated groups. Data are presented as a mean ± standard error of the mean. Differences were tested by Kruskal– Wallis test followed by multiple comparison Dunn’s test. ** P<0.01.

In the acute DSS experiment, faecal iron concentration increased significantly in all DSS-treated mice. This was more pronounced in the 400ppm iron group (Fig. 3-b).

**Figure 3-b:**
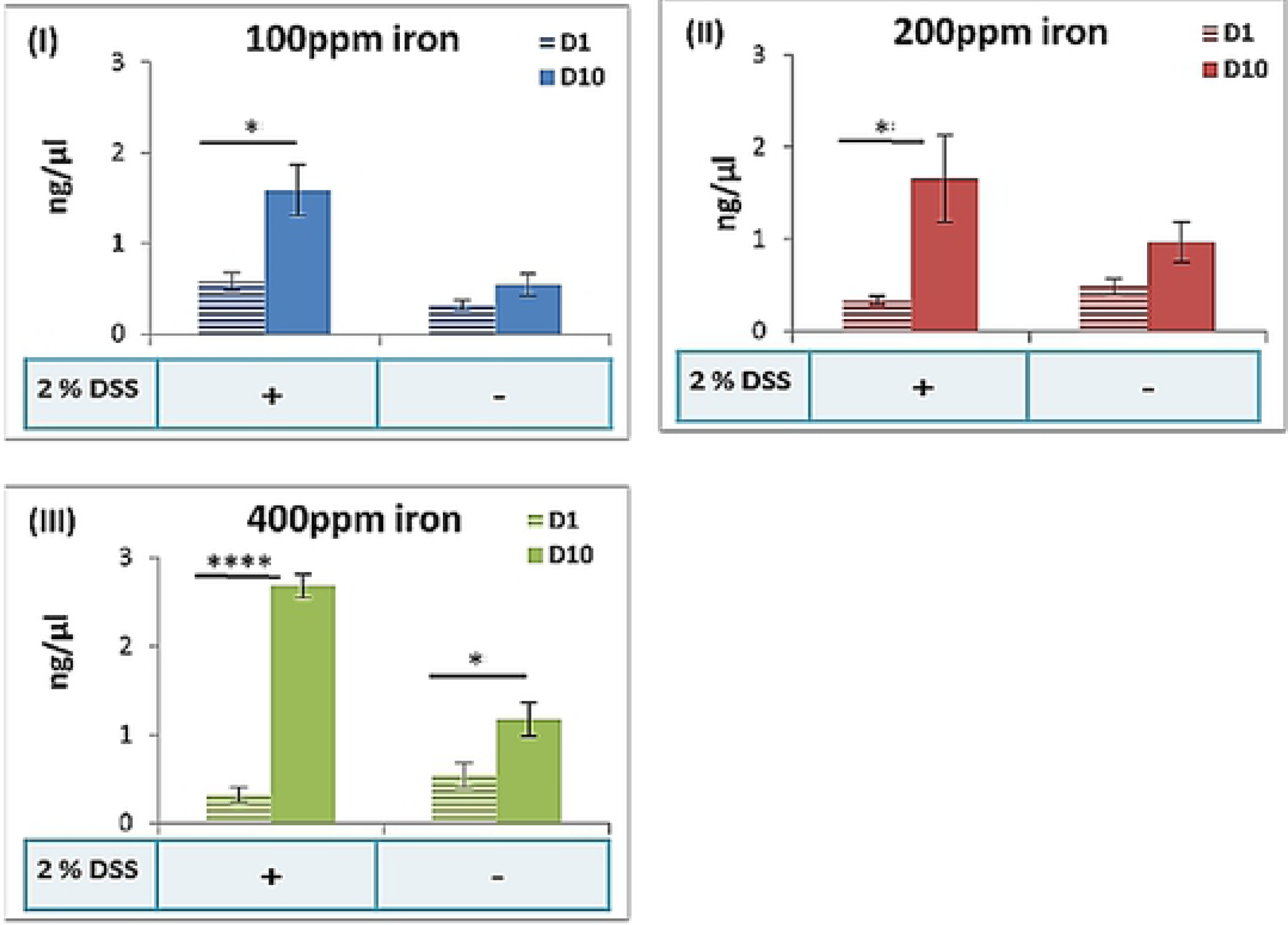
Faecal iron concentration at two different time points day-1 and 10 separately. (I) Faecal iron in 100ppm iron DSS-treated and untreated groups (II) faecal iron in 200ppm iron DSS-treated and untreated groups (III) faecal iron in 400ppm iron DSS-treated and untreated groups. Data are presented as a mean ± standard error of the mean. Differences were tested by Kruskal– Wallis test followed by multiple comparison Dunn’s test.

### Bacterial diversity data analysis at phylum and family level for chronic experiments

Tables of rarefied OTU data were prepared, and three measures of alpha diversity were estimated: chao1, the observed number of species, and the phylogenetic distance. These estimates were plotted as rarefaction curves using Qiime (Supplementary Figure 7). Similarly, for beta-diversity, weighted and unweighted pair-wise UniFrac matrices UPGMA trees were prepared (Supplementary Figure 8).

Principal component analysis (PCA) was used to identify linear combinations of gut microbial taxa associated with the duration on a diet (Fig. 4). Our data showed an overlap in the samples of 100 and 200ppm iron DSS-untreated and 200ppm iron DSS-treated mice (Figure 4-a, c and d). There was clustering with little separation of samples pre- and post-DSS treatment for 100 and the 400ppm iron DSS-treated groups as well as with control mice fed a 400ppm iron diet (Figure 4-b, e and f). The double standard (400ppm) iron diet disturbed the microbial community significantly in both DSS-treated and untreated mice.

**Figure 4:**
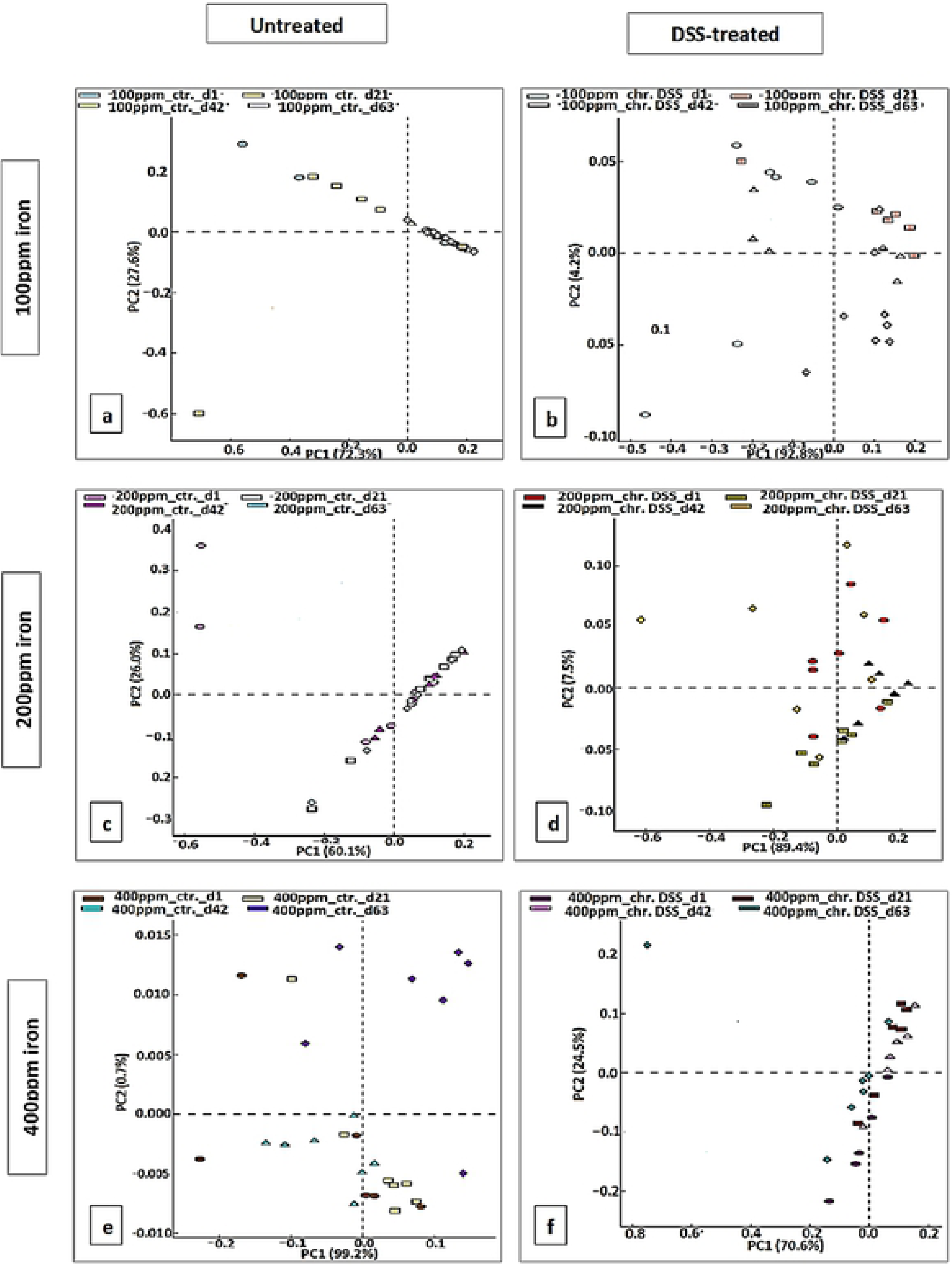
In chronic DSS, PCA plots of the unweighted UniFrac distances of pre-and post-DSS-intervention stool samples from chronic (3 cycles) DSS-treated mice (b, d, and f) and (a, c and e) untreated mice at Phylum-level, phylogenetic classification of 16S rRNA gene sequences. Symbols represent data from individual mice, colour-coded by the indicated metadata. Statistical differences were assessed by Kruskal-Wallis H-test followed by Storey’s FDR multiple test correction.

Post-hoc tests revealed a significant difference in the amount of *Proteobacteria* in 100ppm iron chronic DSS-treated mice when day-1 and 63 were compared (P<0.017) (Fig. 5-a). In 400ppm iron DSS-untreated mice there was a significant increase in two phyla (*Proteobacteria* and *Actinobacteria*) comparing day-1, 21, 42 and 63 samples (p<0.011 for both) (Fig. 5-b). The analysis of faecal samples from mice in the 400ppm iron DSS-treated group showed differences in *Bacteroidetes* and *Proteobacteria* comparing day-1, 21, 42 and 63: *Proteobacteria* increased significantly (P<0.016), and *Bacteroidetes* decreased (P<0.028) (Figure 5-c). Together these data suggest that *Proteobacteria* are dependent on luminal iron, but *Bacteroidetes* are suppressed by inflammation and/or luminal iron.

**Figure 5-a:**
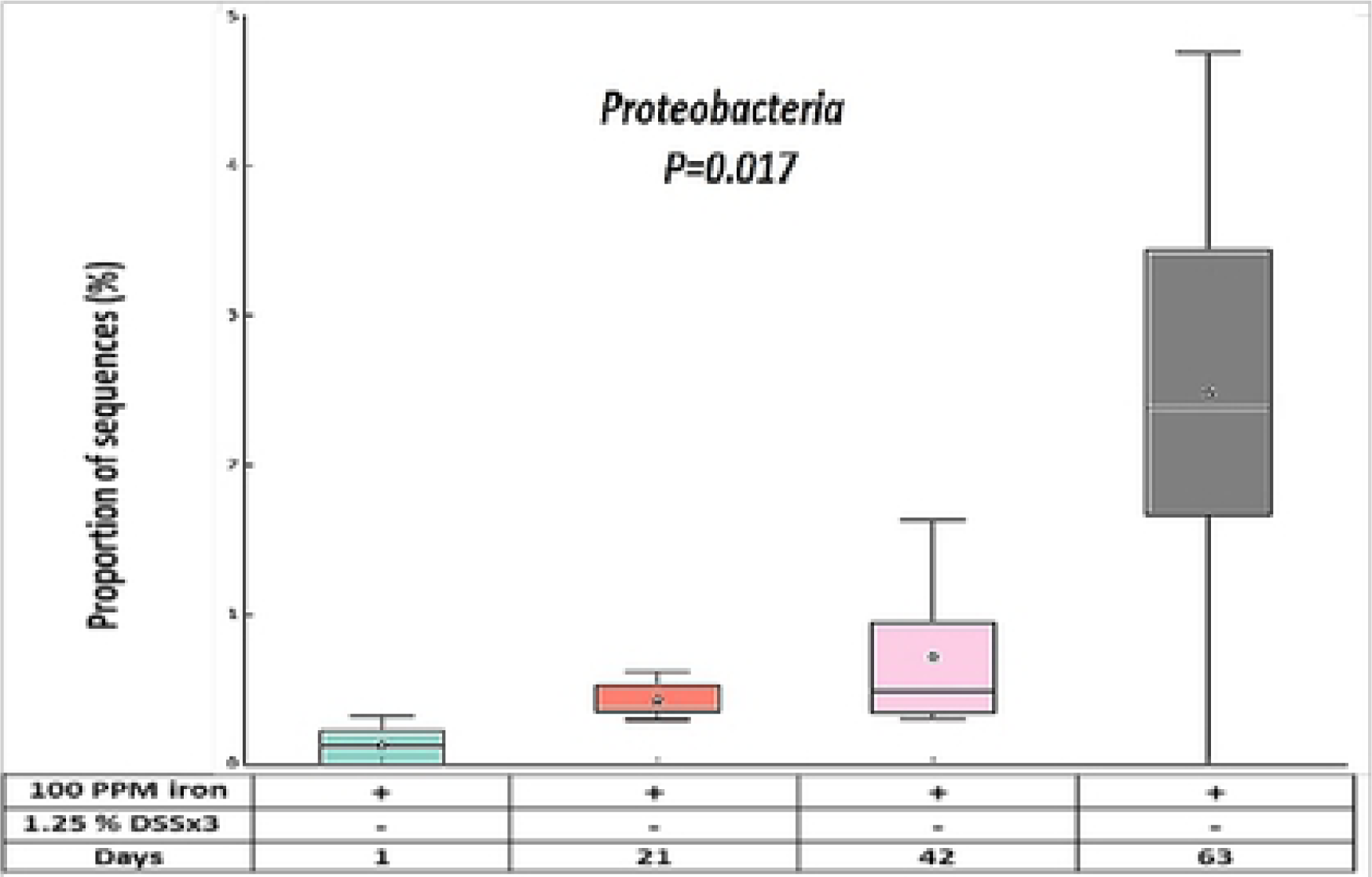
In chronic DSS, box plot showing the distribution in the proportion of *Proteobacteria* assigned to samples at day-1, 21, 42 and 63 from 100ppm iron DSS-treated mice.

**Figure 5-b:**
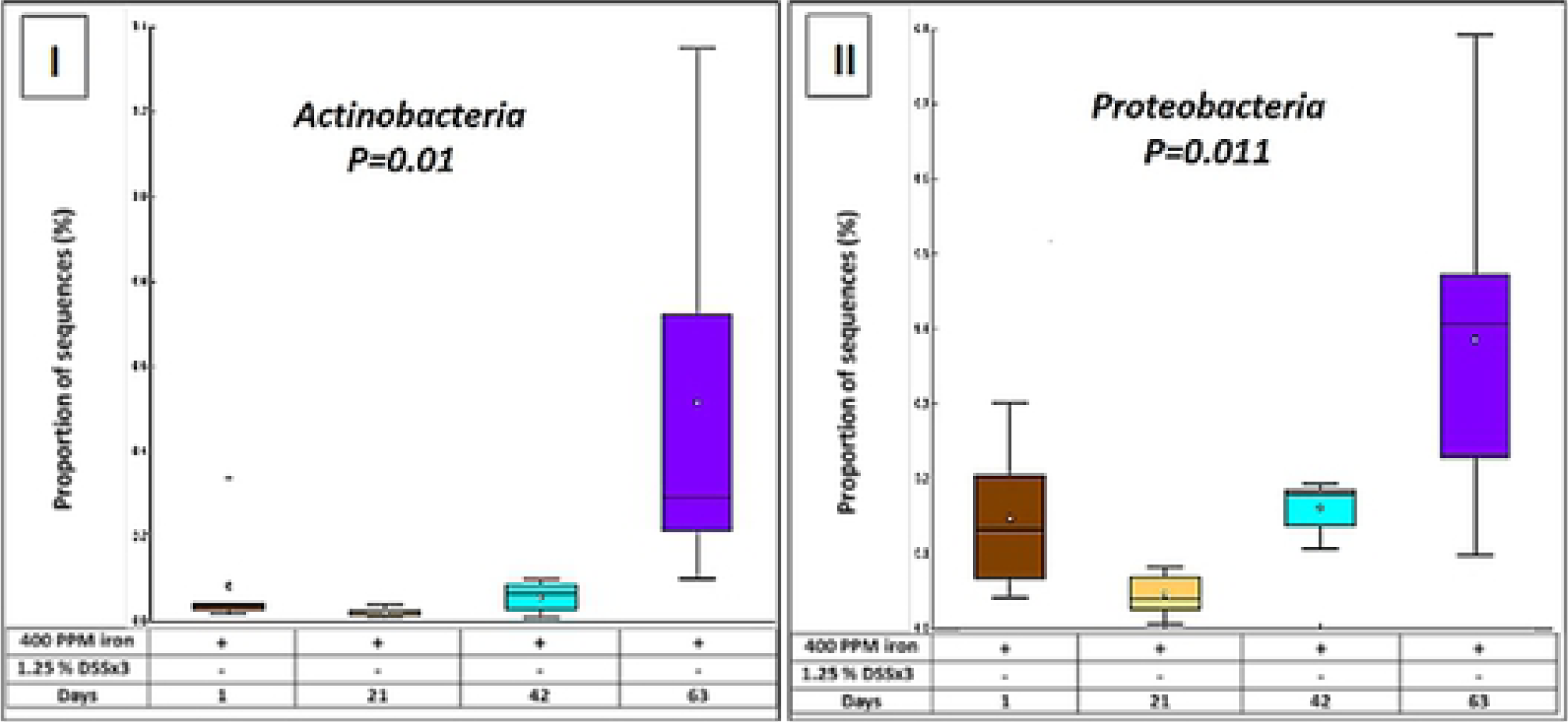
In chronic DSS, box plot showing the distribution in the proportion of two phyla (*Actinobacteria* (I) and *Proteobacteria* (II)) assigned to samples from 400ppm iron untreated mice.

**Figure 5-c:**
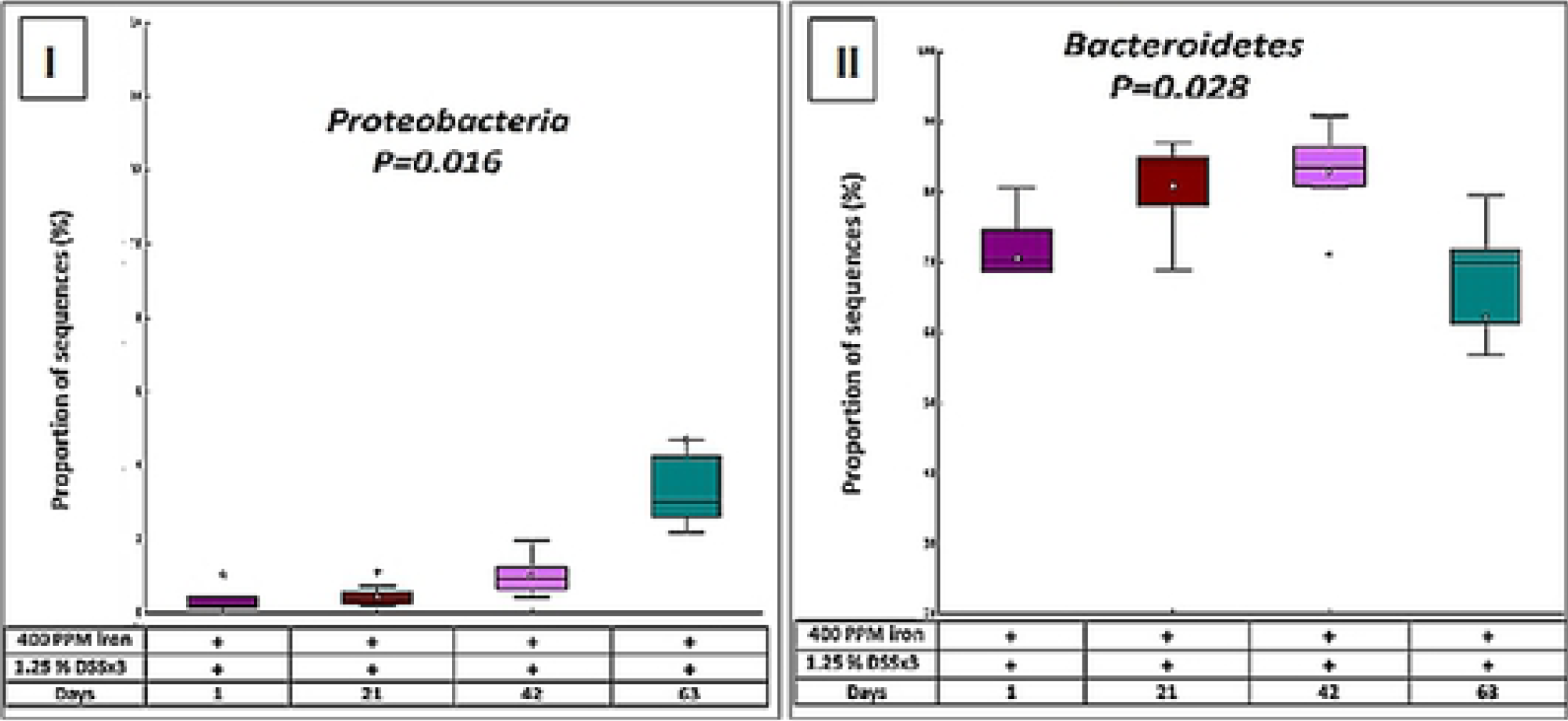
In chronic DSS, box plot showing the distribution in the proportion of two phyla (*Proteobacteria* (I) and *Bacteroidetes* (II)) assigned to samples from 400ppm iron DSS-treated mice.

Further bioinformatics analysis identified 4 phyla and 15 taxa (genera) of interest. Of the four phyla (*Firmicutes, Bacteroidetes, Proteobacteria*, and *Actinobacteria*), one (*Firmicutes*) was highly abundant among all groups while the lowest abundance phylum was *Actinobacteria*. However, 100ppm iron and 400ppm iron chronic DSS groups showed seven different genera apart from the three genera (*Bacteroides, Lactobacillus* and *Bilophila*) that they shared. STAMP encourages the use of effect sizes and confidence intervals (29). The results of the relative abundances of various phyla and identified genera are summarised in Table 1: a-c.

**Table 1-a:** Genus-level taxonomic composition of faecal samples from 100ppm iron DSS-treated mice (Day-1 vs 21, 42 and 63 samples)

| 100ppm iron DSS-treated group |  |  |  |
| --- | --- | --- | --- |
| Taxon | p-values | p-values (corrected) | Effect size |
| p_ *Bacteroidetes*; g_ *Bacteroides* | 0.003 | 0.047 | 0.496 |
| p_ *Bacteroidetes*; g_ *Odoribacter* | 0.002 | 0.04 | 0.620 |
| p_ *Bacteroidetes*; g_ *Prevotella* | 0.0002 | 0.008 | 0.669 |
| p_ *Firmicutes*; g_ *Clostridium* | 0.002 | 0.04 | 0.431 |
| p_ *Firmicutes*; g_ *Dorea* | 0.003 | 0.047 | 0.138 |
| p_ *Firmicutes*; g_ *Lactobacillus* | 0.00002 | 0.002 | 0.880 |
| p_ *Proteobacteria*; g_ *Bilophila* | 0.0002 | 0.008 | 0.766 |

**Table 1-b:** Genus-level taxonomic composition of faecal samples from 400ppm iron DSS-treated mice (Day-1 vs 21, 42 and 63 samples)

| 400ppm iron DSS-treated group |  |  |  |
| --- | --- | --- | --- |
| Taxon | p-values | p-values (corrected) | Effect size |
| p_Firmicutes; g_Lactobacillus | 0.0001 | 0.01 | 0.74 |

**Table 1-c:** Genus-level taxonomic composition of faecal samples from 400ppm iron untreated mice (Day-1 vs 21, 42 and 63 samples)

| 400ppm iron untreated group<br>(Controls) |  |  |  |
| --- | --- | --- | --- |
| Taxon | p-values | p-values (corrected) | Effect size |
| p_Actinobacteria; g_Adlercreutzia | 0.002 | 0.04 | 0.49 |
| p_Bacteroidetes; g_Bacteroides | 0.0005 | 0.02 | 0.68 |
| p_Firmicutes; g_Candidatus Arthromitus | 0.003 | 0.04 | 0.54 |
| p_Firmicutes; g_Lactobacillus | 0.0002 | 0.02 | 0.77 |
| p_Firmicutes; g_Oscillospira | 0.001 | 0.03 | 0.61 |
| p_Firmicutes; g_Ruminococcus | 0.002 | 0.04 | 0.46 |
| p_Proteobacteria; g_Bilophila | 0.001 | 0.03 | 0.55 |

## Discussion

DSS-induced colitis in mice is a popular model for the study of human ulcerative colitis: its mechanism of action is unclear but may be toxic to the colonic epithelium, activate macrophages and/or alter the gut microbiota (30), (31). Most research has used the acute colitis model, however Okayasu *et al* described a chronic colitis model in mice, which may be more appropriate for research of chronic IBD in humans (31) (32) (33). Most studies of the role of iron in relapse of IBD have focussed on the effect of supplementation, however we have recently reported the effect of half standard and double standard dietary iron on acute DSS induced colitis: both changes were associated with more severe colitis than the standard diet (25). Here, we report the effects of the same dietary modification on a model of (1) chronic colitis and (2) acute colitis, in the setting of chronic prior modification of the diet.

When acute colitis was induced after 7 weeks of dietary modification, mice consuming the 100ppm or 400ppm diet developed more severe colitis than mice taking the 200ppm iron diet: clinical and histological data were concordant for 100ppm iron group. In contrast, mice in which chronic colitis was induced while consuming 100ppm, 200ppm or 400ppm dietary iron showed only modest, non-significant weight loss and histological colitis.

In this study, increasing dietary iron led to an increase in faecal iron in the 200 and 400ppm treated mice. After induction of chronic colitis, faecal iron increased in all mice. In the acute DSS experiment, the 400ppm iron group showed the most significant difference (P<0.0001) in faecal iron concentration. There is an obvious paradox: reducing dietary iron was associated with an increase in loss of iron in faeces. The mechanism appears to be by exacerbating DSS-colitis. We speculate that the low iron diet led to more severe colitis, which secondarily led to an increase in bleeding and hence faecal iron.

Changing dietary iron concentration led to a significant difference in the microbiome in both the 100 and 400ppm iron chronic DSS-treated and 400ppm iron untreated groups of animals. Previous research has established a reduction in the biodiversity of commensal bacteria in IBD (34). In mouse experiments, changes in bacterial composition resulted from colonic inflammation and infection (35). In particular, intestinal pathogens (some types of *Proteobacteria*) appeared to take advantage of this. This observation is in agreement with the ‘food hypothesis’ and ‘differential killing’ hypothesis. These two mechanisms are likely to contribute to the loss of colonisation resistance in the inflamed gut (36). Nonetheless, the post-hoc analysis of our data revealed that one bacterial phylum (*Proteobacteria)* was increased significantly (P<0.01) in the 100ppm iron and 400ppm iron DSS-treated and 400ppm iron untreated groups. *Bacteroidetes* decreased significantly (P<0.028) in the 400ppm iron DSS-treated group.

Haller et al. (37) investigated the effects of dietary iron upon the microbiome. Eight bacterial families and nine bacterial genera were significantly (P<0.01) affected by luminal iron (ferrous sulphate) deficiency. The genera *Bifidobacterium* (P<0.0018), *Succinivibrio* (P<0.0027), *Turicibacter* (P<0.0020) and *Clostridium* (P<0.0017) were significantly increased in mice fed an iron depleted diet, whereas the genera *Desulfovibrio* (P<0.0001), *Dorea* (P<0.01) and *Bacteroides* related were greatly reduced. The authors concluded that all significant differences in bacterial abundance in wild-type mice appeared as a result of the interaction between treatment and host-mediated inflammation (37, 38). There are several key differences between that paper and our own: they investigated caecal contents, not faeces; they induced ileitis, not colitis and they did not measure faecal iron concentration. Thus, their paper and our data cannot be directly compared.

Our data analysis showed that seven genera were significantly different. In the half standard iron diets (100ppm) DSS-treated group, we found reductions in *Lactobacillus* (P<0.002), *Dorea, Clostridium, Bacteroides* and *Odoribacter* (P<0.04), *Bilophila* (P<0.008), and an increase in (*Prevotella* P<0.008), all belonging to three phyla [*Firmicutes, Bacteroidetes* and *Proteobacteria*]. In the 400ppm iron DSS group, a significant reduction was shown in *Lactobacillus* (P<0.01). The only control group in which significant differences were found was the 400ppm iron group, where four phyla [*Firmicutes, Bacteroidetes, Proteobacteria*, and *Actinobacteria*] with seven genera showed statistically significant differences. Increases were shown in *Lactobacillus* (P<0.02), *Oscillospira* (P<0.03), *Adlercreutzia* and *Candidatus Arthromitus* (P<0.04), whereas reductions occurred in *Bacteroides* (P<0.02), *Bilophila* (P<0.03) and *Ruminococcus* (P<0.04) (Table 1-c).

Dietary iron plays a role in modulating the susceptibility to DSS-induced colitis. Lower (half standard) iron content in the diet significantly worsened acute colitis leading to an increase in faecal iron. Double standard iron diets caused a dysbiosis. These observations demonstrated the importance of luminal iron and inflammation. Manipulations in dietary iron administration for a longer period significantly exacerbated susceptibility towards developing DSS-induced intestinal inflammation suggesting that the time of iron supplementation may be crucial in aggravating colitis. We cannot explain why the reduced iron diet exacerbates colitis. Further studies will be necessary to investigate the relevance of our findings in humans.

## Supplementary

All data files uploaded in supporting information file.

## “Authors’ contributions.”

**Awad Mahalhal** [conceived and designed research, performed experiments, analysed data, interpreted results of experiments, prepared figures, drafted manuscript, edited and revised manuscript and approved the final version of manuscript]

**Michael D. Burkitt** [analysed data, edited and revised manuscript, approved the final version of manuscript]

**Carrie A. Duckworth** [conceived and designed research, edited and revised manuscript, approved the final version of the manuscript]

**Georgina L. Hold** [analysed data, prepared figures, drafted manuscript, edited and revised manuscript, approved the final version of manuscript]

**Barry J. Campbell** [conceived and designed research, interpreted results of experiments, edited and revised manuscript, approved the final version of manuscript]

**D. Mark Pritchard** [conceived and designed research, edited and revised manuscript, approved the final version of the manuscript

**Chris S. Probert** [conceived and designed research, analysed data, interpreted results of experiments, edited and revised manuscript, approved the final version of manuscript].

